# Different alleles of the same gene vary in pleiotropy, often mediated through currency metabolite production

**DOI:** 10.1101/144329

**Authors:** Deya Alzoubi, Abdelmoneim Amer Desouki, Martin J. Lercher

## Abstract

A major obstacle to the mapping of genotype-phenotype relationships is pleiotropy, the tendency of mutations to affect seemingly unrelated traits. Pleiotropy has major implications for evolution, development, ageing, and disease. Except for disease data, pleiotropy is almost exclusively estimated from full gene knockouts. However, most deleterious alleles segregating in natural populations do not fully abolish gene function, and the degree to which a polymorphism reduces protein function may influence the number of traits it affects. Utilizing genome-scale metabolic models for *Escherichia coli* and the baker’s yeast *Saccharomyces cerevisiae*, we show that most fitness-reducing full gene knockouts of metabolic genes have pleiotropic effects, *i.e.*, they compromise the production of multiple biomass components. Alleles of the same gene with increasingly reduced enzyme function typically affect an increasing number of biomass components. This increasing pleiotropy is often mediated through effects on the generation of currency metabolites such as ATP or NADPH. We conclude that the physiological effects observed in full gene knockouts will in most cases not be representative for alleles with only partially reduced enzyme capacity or expression level.

## Significance statement

Mutations often affect multiple, seemingly unrelated traits, a phenomenon termed pleiotropy. While most mutated genes found in natural populations still function at least partially, experiments on pleiotropy typically knock out genes completely. Using mathematical models for the metabolism of *E. coli* and the baker’s yeast, we show that gene pleiotropy is widespread and modular. Pleiotropy is in many cases mediated through compromised production of important metabolites such as the “energy currency” ATP. Small-effect mutations typically affect only a few traits, while increasingly devastating mutations affect increasing numbers of traits. Thus, full gene knock-outs may represent the pleiotropy found in the natural world, and may be of limited utility for understanding genetic disease and aging, two areas strongly affected by pleiotropy.

## Introduction

A gene is pleiotropic if it affects more than one phenotypic trait (1, 2). A classic example is phenylketonuria, a human disease that is caused by a single gene defect but which affects multiple systems, with symptoms ranging from lighter skin color to mental disorders (3). Pleiotropic effects can cause alleles to affect fitness differentially at different ages, a phenomenon believed to be a major cause of aging (4-6); indeed, alleles contributing to increased longevity often show reduced fertility and stress tolerance (e.g., (7)). Similar antagonistic epistasis may underlie other important biological phenomena such as speciation (8) and cooperation (9). Understanding the factors that contribute to pleiotropy is of fundamental importance in genetics (10-12), evolution (13-16), development (17, 18), as well as in disease (19, 20) and ageing (4). In comparison to its fundamental importance, empirical knowledge of the prevalence and especially on the causal mechanisms of pleiotropy is scarce (2, 21).

Experimental studies generally assess pleiotropy through the analysis of gene knockouts (22-24). The degree of pleiotropy is then defined as the number of traits affected when a gene becomes fully non-functional. Wang *et al*. (24) analyzed phenotypes of large numbers of yeast, nematode, and mouse mutants. They found that pleiotropy is widespread: on average, yeast gene knockouts affect 8% of the examined traits; for the nematode, the corresponding number is 10%, for the mouse 3% (see also (23)). The distributions of the degree of pleiotropy appear rather similar across very different study systems, from the skeletal features of mice (24) to metabolic systems (25-27). Moreover, pleiotropy was found to be modular, such that sets of genes tend to affect the same sets of traits (24).

In *E. coli*, 36% of metabolic reactions are catalyzed by enzymes also involved in other reactions; the same is true for 27% of metabolic reactions in the yeast *Saccharomyces cerevisiae* (27). Pleiotropic effects of mutations that affect enzyme activity can be simulated from genome-scale metabolic models using constraint-based modeling techniques such as flux balance analysis (FBA) (28, 29). The functional pleiotropy of a metabolic gene can then be defined as the number of biomass components whose maximal production is affected by the gene’s knockout (25). Previous studies using this definition found that a metabolic gene’s functional pleiotropy is related to its propensity to form negative epistatic interactions with other metabolic genes (25, 26).

While full gene knockouts are easily examined experimentally, they may not be representative of the effects of deleterious alleles segregating in natural populations: individual mutations may affect only a subset of all traits influenced by the gene (30). Thus, it is important to distinguish between the pleiotropy of the gene and the pleiotropy of individual mutations, especially in evolutionary and clinical contexts. For example, while 4.6% of human SNPs implicated in complex non-Mendelian phenotypes show pleiotropic effects, most of these do not fully abolish protein function (31). Experimental studies indicate that mutational pleiotropy tends to be smaller than gene pleiotropy (30).

Genome-scale metabolic models allow us to dissect the relationship between gene and mutational pleiotropy in quantitative detail, without being hampered by the detection limits of experimental assays. Does the degree of pleiotropy depend on how severely a given allele of a metabolic gene reduces protein activity, *i.e.*, are the same number of functions affected when protein function or expression is reduced only partially? To what extent is pleiotropy mediated through currency metabolites, such as ATP and NADPH, where a reduced production capacity may affect large numbers of otherwise unrelated processes? How modular is metabolic pleiotropy? Below, we address these questions by analyzing the metabolic networks of a representative bacterial model system, *Escherichia coli*, and a corresponding eukaryotic system, the baker’s yeast *Saccharomyces cerevisiae*. We find that most gene knockouts that impact fitness do so by affecting the production of multiple biomass components, and that the number of affected biomass components typically increases with increasing mutation severity. Pleiotropy is rarely a consequence of multiple molecular gene functions, but is an emergent property of the metabolic network; for many genes, pleiotropy is mediated through their involvement in the generation of currency metabolites.

## Results and Discussion

### Estimating pleiotropy from contributions to biomass components within the wildtype flux distribution

We first estimated wildtype flux distributions in the default growth condition for the genome-scale metabolic model of *E. coli* (32) and the yeast *S. cerevisiae* (33) (obtained from https://sourceforge.net/projects/yeast/files/). The maximal biomass production rates were estimated using flux balance analysis (FBA) (28, 29). For both model systems, we identified the flux distribution compatible with maximal biomass production that had the smallest sum of absolute fluxes, a strategy often termed parsimonious FBA (pFBA), which approximates optimal utilization of limited cellular protein resources (34).

To simulate mutations that cause different reductions of protein function or expression and correspond to different deleterious alleles of a metabolic gene, we restricted the maximal flux through all reactions requiring this gene to a fixed percentage of the estimated wildtype flux (35), starting from 100% (the wildtype) down to 0% (a full gene knockout) in steps of 0.5%. For each flux reduction, we defined the degree of pleiotropy (referred to simply as “pleiotropy” below) as the number of biomass components whose production was reduced compared to the maximal (wildtype) production. Note that with this definition, only genes with pleiotropy ≥2 are pleiotropic, while genes with pleiotropy 0 (no affected biomass component) or pleiotropy 1 (one affected biomass component) are non-pleiotropic.

Flux distributions at maximal biomass production rate are usually not unique (34), and so in many cases a flux restriction through one reaction may be compensated by a rerouting of fluxes through alternative pathways. However, such rerouting would require the upregulation of the corresponding genes, which is unlikely to occur spontaneously (36). More importantly, if we are interested in the *de facto* contribution of a given gene to the production of biomass components, then it is of no consequence if alternative pathways *could* take over part of this functionality. Thus, when calculating the maximal (wildtype) production rate of individual biomass components as well as when simulating the effects of mutations to a given metabolic gene, we did not allow the redistribution of fluxes to alternative pathways: we allowed only decreases, not increases, of any fluxes compared to the wildtype flux distribution obtained with pFBA.

### Many genes affect the production of multiple biomass components

Pleiotropy varies widely between different genes. Mutations to the majority of genes affect no biomass components in the minimal growth medium assayed, independent of mutation severity (*E. coli*: 78.13% of genes; *S. cerevisiae*: 75.58%). Among genes contributing to biomass production—and thus fitness—in the wildtype, non-pleiotropic cases are rare: in *E. coli*, only 54 full-gene knockouts (out of 299 knockouts with fitness contributions) affect exactly one biomass component, while the same is true for only 12 knockouts (out of 222 knockouts with fitness contributions) in *S. cerevisiae*. Conversely, knockouts of 32 genes in *E. coli* and 40 genes in *S. cerevisiae* affect the production of *all* biomass components. Many of the remaining genes show low degrees of pleiotropy, affecting the production of only a few biomass components; on average, full gene knockouts of fitness-relevant genes affect the production of 20% of biomass components in *E. coli* and 34% of biomass components in *S. cerevisiae* (Figure 1, Table 1).

**Figure 1.**
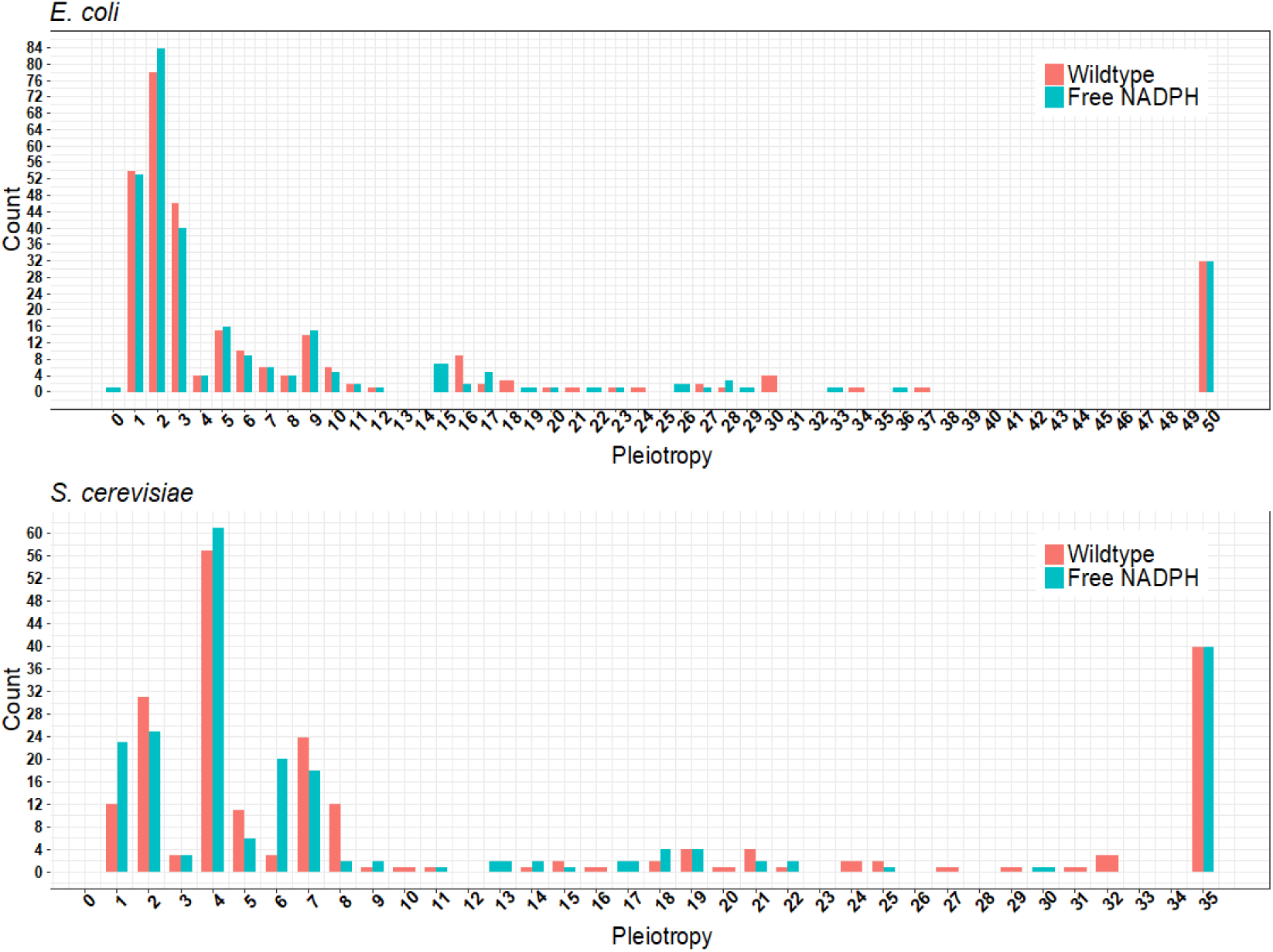
Most complete gene knockouts of fitness-relevant genes have pleiotropic effects, *i.e.*, they affect the production of multiple biomass components. For some genes, pleiotropy is reduced when NADPH is made freely available (green bars). For other freely available currency metabolites, see Supplementary Figure S2.

**Table 1:** Average number of biomass components whose production is affected by a full gene knockout.

|  |  | *E. coli*<sup>3</sup> | *S. cerevisiae*<sup>3</sup> |
| --- | --- | --- | --- |
| # Biomass components |  | 50 | 35 |
| Standard model | Pleiotropy<sup>1</sup> | 10.0 (3) | 12.1 (5) |
|  | Essentiality<sup>1</sup> | 4.6 (2) | 9.3 (2) |
| Free NADPH<sup>2</sup> | Pleiotropy<sup>1</sup> | 9.9 (3) | 11.1 (4.5) |
|  | Essentiality<sup>1</sup> | 4.3 (2) | 8.0 (1) |

<sup>1</sup> In the FBA calculations, fluxes are either constrained to not exceed the wildtype (WT) fluxes to estimate the *de facto* contribution of gene products to biomass production (pleiotropy), or they are allowed to vary freely to assess the number of biomass components for which gene products are essential even after allowing the mutant strain to adapt.
<sup>2</sup> Solution when allowing unlimited conversion of NADPH to NADP+
<sup>3</sup> Mean (median) number of affected biomass components

These percentages reflect functional pleiotropy, the *de facto* contribution of gene products to biomass component production. If, instead, we are interested in the phenotypic effects of gene knockouts after allowing the mutant strain to adapt its physiology to its altered gene content, we must allow free redistributions of fluxes after the gene knockouts. Corresponding simulations show that genes with fitness contributions are, on average, essential for the production of 9.2% of *E. coli* biomass components and of 26.6% of *S. cerevisiae* biomass components after adaptation (Table 1, Supplementary Figure S1). The degree of gene pleiotropy for yeast is higher than previous experimental estimates, which range from 2%-11% depending on the considered traits (24); however, experimental estimates of gene pleiotropy tend be downwardly biased due to experimental detection limits (2, 37).

### Pleiotropy is an emergent property of the metabolic network

Pleiotropy can be classified by its origin into type I pleiotropy, caused by multiple molecular functions of a gene product, and type II pleiotropy, caused by multiple physiological consequences of a single molecular function (2). Similar distinctions have been made previously using the terms “horizontal” vs. “vertical” (10) and “mosaic” vs. “relational” (38) pleiotropy. Our model allows us to quantify the relative contributions of these two pleiotropy types. 41.7% of *E.coli* genes and 40.5% of yeast genes in our metabolic models catalyze multiple reactions. To what extent does this functional diversity cause functional pleiotropy as measured in the number of biomass components affected by a gene knockout? To answer this question, we compared the gene pleiotropy (Fig. 1) to the pleiotropy of individual reactions catalyzed by the gene product. For example, fully abolishing all functions of the purB (b1131) gene, whose gene product catalyzes two distinct biochemical reactions, reduced the production of 18 biomass components. In contrast, blocking only one of the catalyzed reactions results in a pleiotropy estimate of 16, while blocking only the other reaction results in a pleiotropy of 10. Thus, the pleiotropy of the b1131 gene is largely of type II, and is only is only in small part due to its multiple molecular functions.

This pattern is typical: the maximal pleiotropy arising from blocking only a single out of several reactions catalyzed by the same protein accounts for over 97% of the gene pleiotropy (*E. coli* 97.4%, yeast 97.6%). These numbers drop only marginally when we consider only gene products that are essential for multiple reactions, to 92.2% in *E. coli* and to 94.5% in yeast (Supplementary Figure S3). We conclude that the vast majority of metabolic epistasis is of type II, *i.e.*, is an emergent property of the metabolic network rather than a consequence of multiple molecular functions. This finding is consistent with the previous observation that the degree of pleiotropy in yeast is not significantly correlated with the number of molecular gene functions (39).

### Metabolic networks show significant but low modularity

The relationship between genes and biomass components (traits) can be represented as a bipartite graph, with links connecting genes with affected biomass components. Modules are defined as sets of genes and traits with significantly more within-module than between-module links (24). Supplementary Figure S4 shows heatmaps that illustrate the modularity of both metabolic pleiotropy networks. To quantitatively assess the modularity, we used the LP&BRIM algorithm (40), resulting in raw modularities of Q=0.235 for *E. coli* and Q=0.197 for *S. cerevisiae*. Both networks show highly statistically significant modularity: in each case, the modularity of 10,000 randomly rewired networks was always lower that observed for the real pleiotropy network (*i.e., P*<0.0001; Supplementary Figure S5).

Following Ref. (24), we then defined the scaled modularity as the difference between the observed modularity and the mean modularity of randomly rewired networks, measured in number of standard deviations (41). The *E. coli* pleiotropy network exhibits a scaled modularity of 9.1, while the *S. cerevisiae* network has a scaled modularity of 4.9, *i.e.*, the modularity of metabolic pleiotropy is about 9 and 5 standard deviations higher than for corresponding random gene-trait networks. These values are surprisingly low: for five different experimental study systems and trait definitions, Wang *et al.* found a median scaled modularity of 37 (range 34-238).

### Pleiotropy typically increases with increasing mutation severity

We next examined the pleiotropy of alleles with small-effect mutations, *i.e.*, mutations that reduce enzyme capacity without fully abolishing enzyme function. About 20% of *E. coli* genes with fitness contributions have constant pleiotropy: small-effect mutations of these genes affect the same number of biomass components as full gene knockouts. In comparison, only 7.7% of yeast genes contributing to fitness exhibit constant pleiotropy.

All other genes contributing to fitness affect an increasing number of biomass components for increasingly deleterious alleles. Figure 2 shows this stepwise increase in pleiotropy for the example of *Lipoamide dehydrogenase* (gene names: *E. coli* b0116, *S. cerevisiae* YFL018C; for additional examples, see Supplementary Figure S6. In both organisms, pleiotropy typically increases in about a dozen steps from weakly to strongly deleterious alleles (Figure 3; mean number of steps: *E. coli* 11.6, *S. cerevisiae* 12.6).

**Figure 2.**
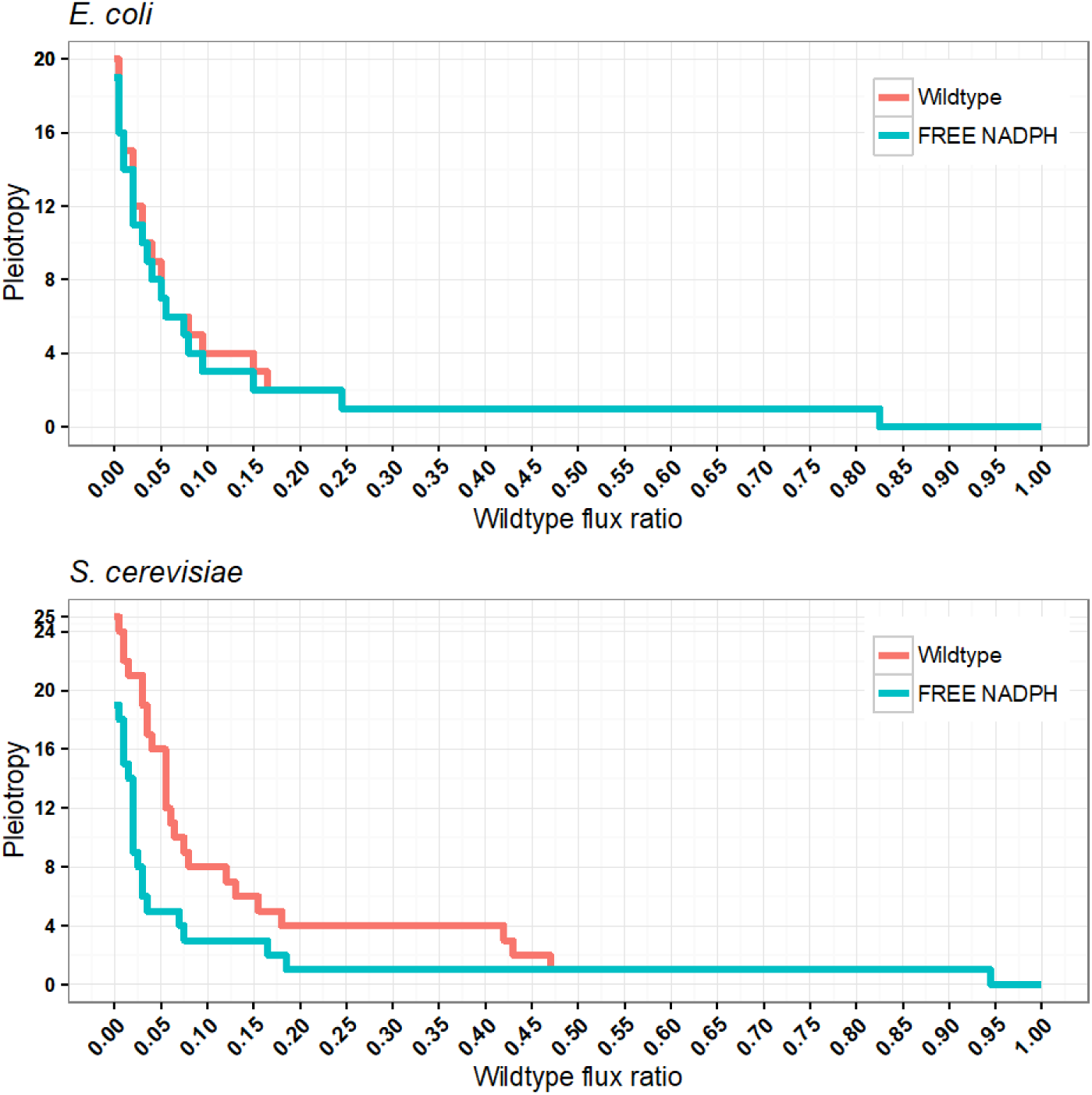
Pleiotropy for the *Lipoamide dehydrogenase* gene increases for increasingly deleterious alleles. Pleiotropy is reduced when NADPH is made freely available (green curves). For additional examples, see Supplementary Figure S4-6X.

**Figure 3.**
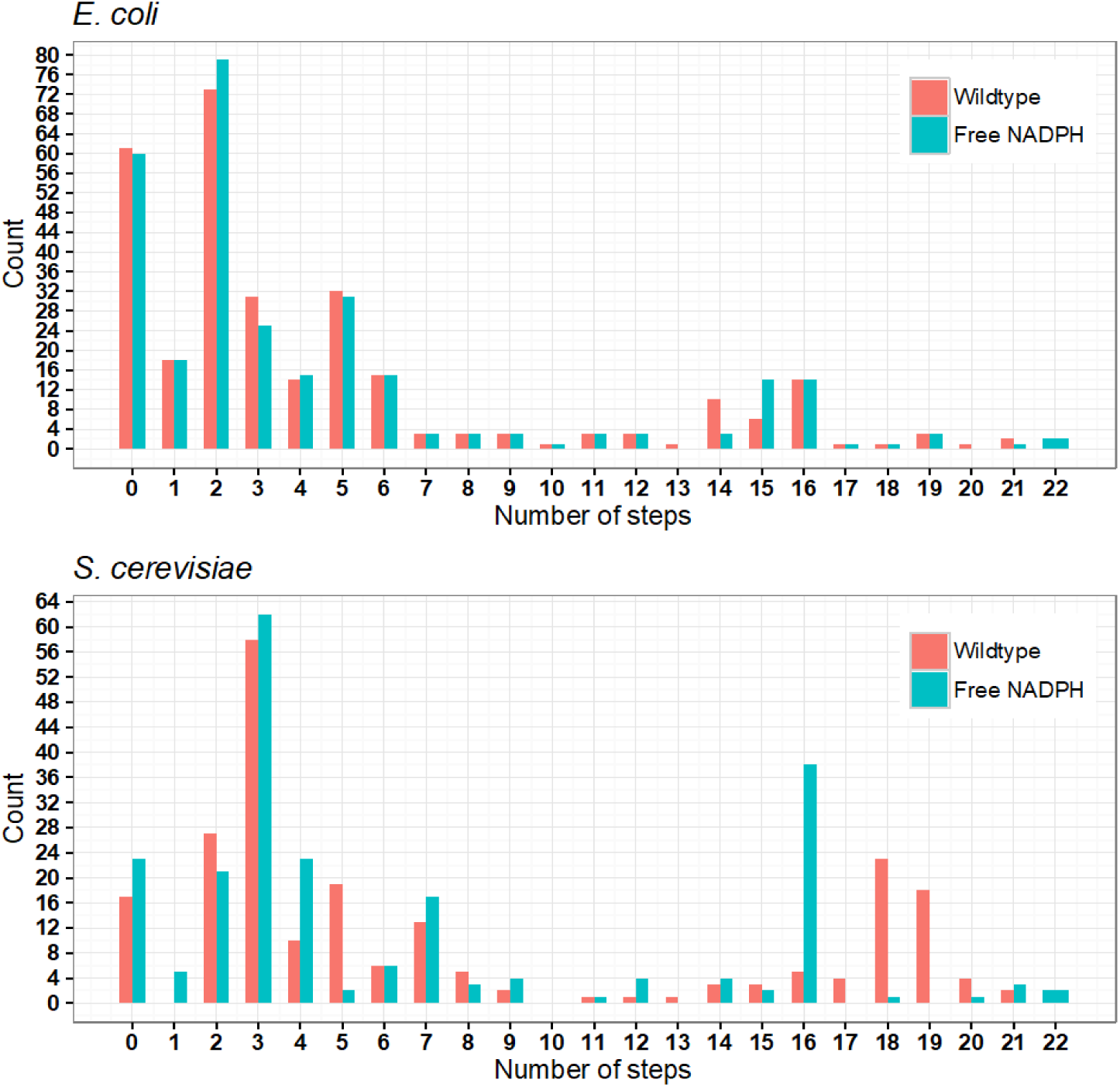
For the majority of genes contributing to biomass production, pleiotropy increases for increasingly deleterious alleles in multiple steps. Histograms for the number of pleiotropy steps in *E. coli* and the yeast *S. cerevisiae*. Green bars reflect the reduced numbers of pleiotropy increases when making NADPH freely available.

The pleiotropy of the full gene knockout constitutes an upper limit to the number of stepwise increases in pleiotropy. However, the correlation between the number of steps and pleiotropy at full knockout was even stronger than expected from this relationship (Supplementary Figure S7, Spearman’s *ρ*=0.926 (*E. coli*) and *ρ*= 0.986 (*S. cerevisiae*), *P*<,10^−15^ in each case). Thus, genes whose full knockout showed higher metabolic pleiotropy also showed more stepwise increases in pleiotropy for increasingly debilitating mutations.

All genes whose mutations affect the production of at least one biomass component must also affect the overall production of biomass (*i.e*., in the common interpretation of FBA, fitness). The reverse is not true: a mutation to a gene may affect the maximal production of biomass, but not the production of any individual biomass component. This is a consequence of the algorithm employed to estimate production capabilities for individual biomass components. If we maximize the production of a single compound, then pathways usually concerned with the production of other biomass components can be diverted to the production of this compound. While we find no such genes for *S. cerevisiae*, this is indeed the case for 3 essential *E. coli* genes, which encode transporters for acetate (b4067), magnesium/nickel/cobalt (b3816), and calcium/sodium (b3196, an antiporter).

### The pleiotropy of most genes is mediated by currency metabolites

We can conceptually partition internal metabolites into currency metabolites—those involved in many reactions, *e.g*., to provide energy or redox equivalents (42)—and primary metabolites. A deleterious allele may affect the production of a given biomass component because the mutated gene catalyzes a reaction in a pathway of primary metabolites that directly leads to the component’s production. Conversely, a deleterious allele may affect not the primary metabolites, but the currency metabolites utilized in the component’s production.

A substantial fraction of pleiotropy is indeed associated with the generation of currency metabolites: 87.4% of previously pleiotropic genes show reduced pleiotropy when we make metabolites such as ATP, UTP, or NADPH freely available in yeast (Figure 4). The free availability of ATP alone reduces the degree of pleiotropy of over half of pleiotropic yeast genes. The influence of currency metabolite production on pleiotropy is weaker, yet still substantial in *E. coli*: here, 55.3% of pleiotropic genes are affected, with NADH making the biggest contribution (over 40%) (Figure 4).

**Figure 4.**
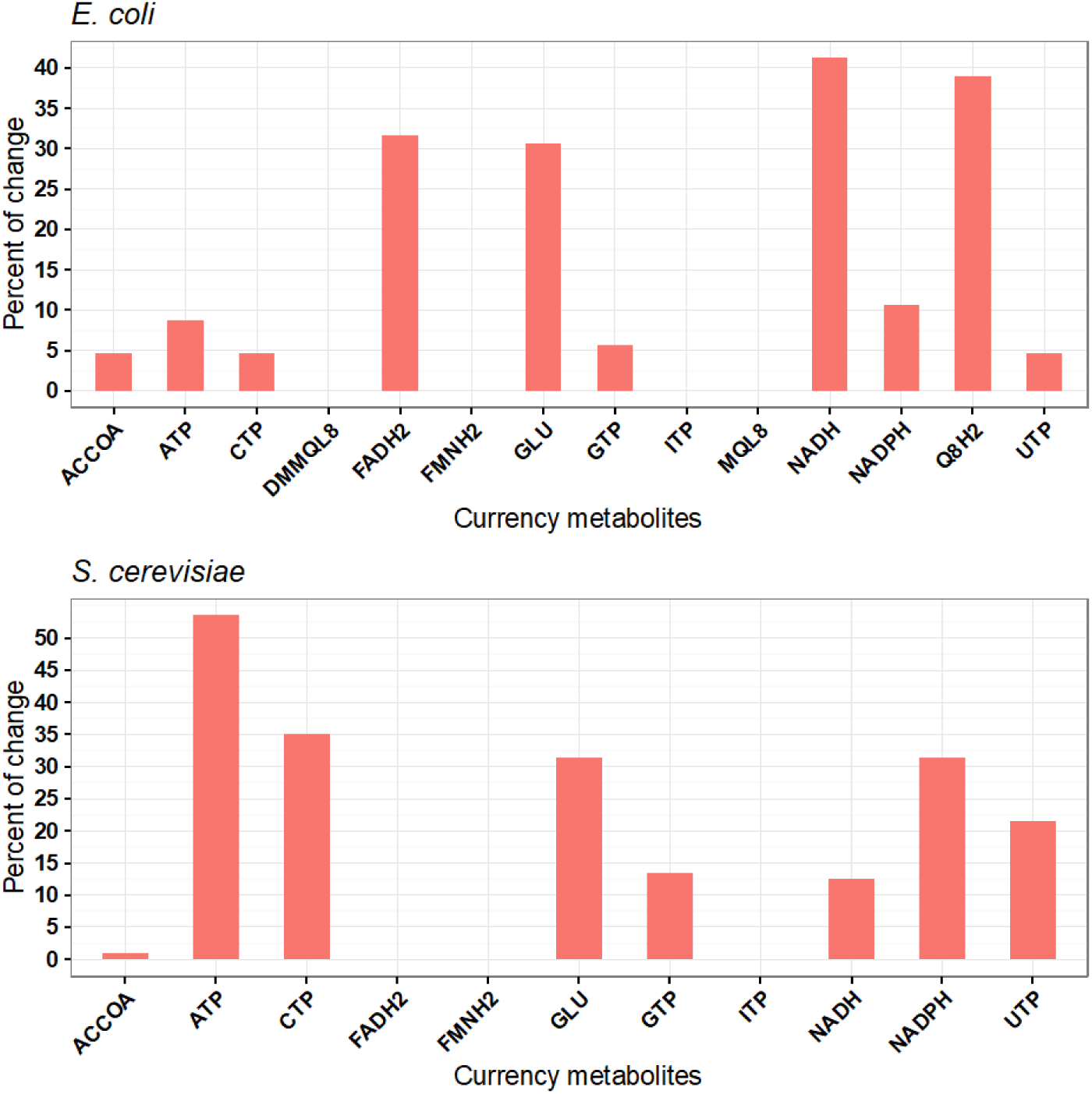
Many genes show reduced pleiotropy when currency metabolites are made freely available. The bar chart shows the percentage of previously pleiotropic genes with reduced pleiotropy in response to the free availability of different currency metabolites. Abbreviations: Adenosine triphosphate (ATP); Cytidine triphosphate (CTP); Guanosine triphosphate (GTP); Uridine triphosphate (UTP); Inosine triphosphate (ITP); Nicotinamide adenine dinucleotide (NADH); Nicotinamide adenine dinucleotide phosphate (NADPH); Flavin adenine dinucleotide reduced (FADH2); Reduced flavin mononucleotide (FMNH2); Ubiquinol-8 (Q8H2); Menaquinol 8 (MQL8); 2-Demethylmenaquinol 8 (DMMQL8); Acetyl-CoA (ACCOA); L-Glutamate (GLU).

Involvement in currency metabolite production is an important determinant of the number of biomass components for which a gene knockout is essential even after allowing the mutant strain to adapt its protein expression to the altered gene content of its genome. This contribution is particularly striking in yeast: for over half of the tested currency metabolites, free availability reduces the number of biomass components for which a gene is essential for almost half of the genes (Supplementary Figure S8).

## Conclusion

Using constraint-based simulations of the metabolic models for *E. coli* and the yeast *S. cerevisiae*, we have characterized the distributions of pleiotropy. Consistent with earlier computational (25-27) and experimental (22-24) studies, we found that the knockout of a majority of genes that contribute to fitness has pleiotropic effects. The vast majority of this gene pleiotropy is not caused by multiple molecular functions of the gene product (type I), but is an emergent property of the metabolic network (type II). Pleiotropy is modular, but to a lower degree than estimated experimentally for non-metabolic systems (24).

For most pleiotropic genes, pleiotropy increases strongly for alleles with increasingly debilitating effects. Thus, standard measures of pleiotropy based on gene knockout studies are more likely to reflect the maximal degree of mutational pleiotropy of a given gene (2, 30). Alleles that only knock down protein activity (by reducing enzyme/transporter function or expression level) often affect only a subset of phenotypic traits, with additional traits affected progressively as alleles become more deleterious. Thus, the physiological effect of the full gene knockout will in most cases not be representative for the effects of deleterious alleles that retain some level of enzyme function. This type of effect is also evident from individual medical observations of pleiotropy. For example, some small-effect mutations affecting human SOX9 expression lead to minor skeletal malformations, while the consequences of large-effect mutations can include sex reversals (43).

How can we understand the dependence of pleiotropy on the degree to which an allele reduces protein activity? For increasingly deleterious alleles, more and more metabolic resources must be channeled into the compensation of the compromised pathway; as a consequence of this increasing drain of resources, more and more other pathways are affected. Perhaps not surprisingly (26), we found that the pleiotropy of many genes is mediated through the generation of currency metabolites such as ATP, NADPH, or FADH_2_. This is true for more than half of the pleiotropic genes in *E. coli*, and for 87% of pleiotropic genes in yeast.

Pleiotropy is a complex phenomenon: it is not constant, but varies between different alleles of the same gene, and its causes are often indirect. Thus, experimental as well as computational analyses of pleiotropy should move away from focusing on full gene knockouts, and instead consider explicitly the degree to which mutations reduce protein activity. The necessity of a corresponding nuanced view of pleiotropy may be particularly evident in studies of medically relevant mutations, where full knockouts are often lethal, while small-effect mutations may segregate at appreciable frequencies in the human population (44).

## Materials and methods

### Metabolic models

To simulate *Escherichia coli* metabolism, we used the metabolic reconstruction iJO1366 (32), encompassing 1,366 metabolic genes associated with 2,251 reactions. For the yeast *S. cerevisiae*, we used the yeast7.6 model (https://sourceforge.net/projects/yeast) (33), accounting for 909 metabolic genes associated with 3,326 reactions.

The models specify growth reactions with 50 essential biomass components in *E. coli* (Supplementary Table S1) and 35 essential biomass components in *S. cerevisiae* (Supplementary Table S2), excluding inorganic ions, H_2_O, and products of the biomass reaction.

### Flux distribution constraints derived from wildtype simulations

In order to approximate the *de facto* contribution of individual metabolic proteins to the production of individual biomass components *in vivo*, we should only consider flux distributions that are contained within realistic flux distributions – *i.e.*, we should only use reactions that are naturally active during growth (biomass production) in the nutritional environment studied, and fluxes should not exceed these wildtype fluxes. We thus first estimate the wildtype flux distribution **v**_^WT^_, by running a flux balance analysis (FBA) with the biomass reaction as the objective function, followed by a minimization of the sum of absolute fluxes at the previously determined maximal biomass production rate (parsimonious FBA (34)).

When simulating the production of individual biomass components, we constrained all fluxes *v*_*i*_ to values between zero and the wildtype flux 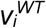 for this reaction, *i.e.*,

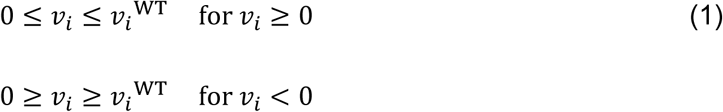

### Estimating pleiotropy

For each essential biomass component, we added a new exchange reaction representing its secretion (45). As some biomass components may be coupled through the biomass reaction, we allowed the excretion of all other biomass components when maximizing the production of one selected biomass component.

We then calculated the maximum production of each biomass component by maximizing its exchange reaction while enforcing the wildtype flux distribution constraints (Eq. 1). For each metabolic gene, we compared this unperturbed production with the maximal production rate of alleles with increasingly reduced protein activity, simulated by restricting the flux through all reactions catalyzed by the gene to a fixed fraction of the wildtype flux (35), which we reduced from 100% to 0% in steps of 0.5%.

We defined pleiotropy as the number of biomass components whose maximal production was reduced compared to the unperturbed state for the allele considered. Thus, an allele not involved in the maximal production of any essential biomass component is considered to have pleiotropy 0; an allele that affects the production of exactly one essential biomass component has pleiotropy 1.

Our estimate of pleiotropy reflects the actual contribution of a gene product to biomass formation, based on estimated enzyme and transporter activities in the wildtype. If instead, one is interested in a quantitative measure of essentiality, defined as the number of biomass components affected by a deleterious allele *after* the mutant strain has been allowed to adapt its physiology to the gene deletion, a different algorithm is more appropriate. In this case, one needs to allow the free redistribution of fluxes after the simulated activity reduction of the protein encoded by the gene in question. Note that experimental studies often employ a pragmatic working definition of pleiotropy that lies somewhere between the definitions of pleiotropy and quantitative essentiality proposed here: in these studies, pleiotropy is typically estimated as the number of traits with observable phenotypic changes after the gene knockout, but before allowing the strain to adapt. In this case, some fluxes may be rerouted due to enzymes and transporters that are expressed either spuriously or because of other roles they play in wildtype physiology, while other fluxes that require the upregulation of the corresponding enzymes and transporters will not yet be active.

### Currency metabolites

In additional analyses, we made several cofactors freely available to study how pleiotropy is associated with the generation of currency metabolites. We did this by adding a balanced biochemical reaction that interconverts the activated and inactivated versions of the cofactor and allowing unlimited flux of this reaction in both directions. For example, to simulate free NADPH, we added the following reversible reaction:

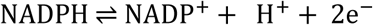

We used the following currency metabolites (46): Adenosine triphosphate (ATP); Cytidine triphosphate (CTP); Guanosine triphosphate (GTP); Uridine triphosphate (UTP); Inosine triphosphate (ITP); Nicotinamide adenine dinucleotide (NADH); Nicotinamide adenine dinucleotide phosphate (NADPH); Flavin adenine dinucleotide reduced (FADH2); Reduced flavin mononucleotide (FMNH2); Ubiquinol-8 (Q8H2); Menaquinol 8 (MQL8); 2-Demethylmenaquinol 8 (DMMQL8); Acetyl-CoA (ACCOA); L-Glutamate (GLU).

### Software used

All simulations were performed in R (47) using *sybil*, a computer library optimized for efficient constraint-based modeling of metabolic networks (48). We used IBM ILOG CPLEX as the linear solver, connected to sybil via the cplexAPI R package.

To calculate network modularities, we used the LP&BRIM algorithm (Label Propagation with Bipartite Recursively Induced Modules) implemented in Matlab (40).

## Acknowledgements

We thank Balazs Papp for helpful discussions. This work was supported financially through fellowships by the German Academic Exchange Service (DAAD) to DA and AAD, and through a German Research Foundation (DFG) grant to MJL in the context of collaborative research centre CRC 680.

## References

1. Stearns FW (2011) One Hundred Years of Pleiotropy: A Retrospective (vol 186, pg 767, 2010). Genetics 187(1):355–355.

2. Wagner GP & Zhang J (2011) The pleiotropic structure of the genotype-phenotype map: the evolvability of complex organisms. Nat Rev Genet 12(3):204–213.

3. Paul D (2000) A double-edged sword. Nature 405(6786):515–515.

4. Williams GC (1957) Pleiotropy, Natural-Selection, and the Evolution of Senescence. Evolution 11(4):398–411.

5. Zwaan BJ (1999) The evolutionary genetics of ageing and longevity. Heredity 82:589–597.

6. Moorad JA & Promislow DEL (2009) What can genetic variation tell us about the evolution of senescence? Proceedings of the Royal Society B-Biological Sciences 276(1665):2271–2278.

7. Paaby AB & Schmidt PS (2009) Dissecting the genetics of longevity in Drosophila melanogaster. Fly (Austin) 3(1):29–38.

8. Slatkin M (1982) Pleiotropy and Parapatric Speciation. Evolution 36(2):263–270.

9. Foster KR, Shaulsky G, Strassmann JE, Queller DC, & Thompson CR (2004) Pleiotropy as a mechanism to stabilize cooperation. Nature 431(7009):693–696.

10. Tyler AL, Asselbergs FW, Williams SM, & Moore JH (2009) Shadows of complexity: what biological networks reveal about epistasis and pleiotropy. Bioessays 31(2):220–227.

11. Wright S (1968) Evolution and the genetics of populations: a treatise in four volumes (University of Chicago Press, Chicago).

12. Barton NH (1990) Pleiotropic Models of Quantitative Variation. Genetics 124(3):773–782.

13. Fisher RA (1930) The genetical theory of natural selection (Clarendon Press, Oxford,) pp xiv, 272 p.

14. Orr HA (2000) Adaptation and the cost of complexity. Evolution 54(1):13–20.

15. Waxman D & Peck JR (1998) Pleiotropy and the preservation of perfection. Science 279(5354):1210–1213.

16. Otto SP (2004) Two steps forward, one step back: the pleiotropic effects of favoured alleles. Proc Biol Sci 271(1540):705–714.

17. Hodgkin J (1998) Seven types of pleiotropy. Int J Dev Biol 42(3):501–505.

18. Carroll SB (2008) Evo-devo and an expanding evolutionary synthesis: a genetic theory of morphological evolution. Cell 134(1):25–36.

19. Albin RL (1993) Antagonistic pleiotropy, mutation accumulation, and human genetic disease. Genetica 91(1-3):279–286.

20. Brunner HG & van Driel MA (2004) From syndrome families to functional genomics. Nat Rev Genet 5(7):545–551.

21. Zhang J & Wagner GP (2013) On the definition and measurement of pleiotropy. Trends Genet 29(7):383–384.

22. Dudley AM, Janse DM, Tanay A, Shamir R, & Church GM (2005) A global view of pleiotropy and phenotypically derived gene function in yeast. Molecular Systems Biology

23. Wagner GP, et al. (2008) Pleiotropic scaling of gene effects and the ‘cost of complexity’. Nature 452(7186):470–472.

24. Wang Z, Liao BY, & Zhang JZ (2010) Genomic patterns of pleiotropy and the evolution of complexity. Proceedings of the National Academy of Sciences of the United States of America 107(42):18034–18039.

25. Szappanos B, et al. (2011) An integrated approach to characterize genetic interaction networks in yeast metabolism. Nat Genet 43(7):656–662.

26. Bajic D, Moreno-Fenoll C, & Poyatos JF (2014) Rewiring of genetic networks in response to modification of genetic background. Genome Biol Evol 6(12):3267–3280.

27. Wang Z & Zhang JZ (2009) Abundant Indispensable Redundancies in Cellular Metabolic Networks. Genome Biology and Evolution 1:23–33.

28. Watson MR (1984) Metabolic Maps for the Apple-II. Biochem Soc T 12(6):1093–1094.

29. Orth JD, Thiele I, & Palsson BO (2010) What is flux balance analysis? Nature Biotechnology 28(3):245–248.

30. Stern DL (2000) Evolutionary developmental biology and the problem of variation. Evolution 54(4):1079–1091.

31. Sivakumaran S, et al. (2011) Abundant Pleiotropy in Human Complex Diseases and Traits. American Journal of Human Genetics 89(5):607–618.

32. Orth JD, et al. (2011) A comprehensive genome-scale reconstruction of Escherichia coli metabolism--2011. Mol Syst Biol 7:535.

33. Aung HW, Henry SA, & Walker LP (2013) Revising the Representation of Fatty Acid, Glycerolipid, and Glycerophospholipid Metabolism in the Consensus Model of Yeast Metabolism. Ind Biotechnol (New Rochelle N Y) 9(4):215–228.

34. Holzhutter HG (2004) The principle of flux minimization and its application to estimate stationary fluxes in metabolic networks. European Journal of Biochemistry 271(14):2905–2922.

35. Xu L, Barker B, & Gu ZL (2012) Dynamic epistasis for different alleles of the same gene. Proceedings of the National Academy of Sciences of the United States of America 109(26):10420–10425.

36. Shlomi T, Berkman O, & Ruppin E (2005) Regulatory on/off minimization of metabolic flux changes after genetic perturbations. Proc Natl Acad Sci U S A 102(21):7695–7700.

37. Paaby AB & Rockman MV (2013) The many faces of pleiotropy. Trends Genet 29(2):66–73.

38. Hadorn E (1961) Developmental genetics and lethal factors (Methuen; Wiley, London, New York,) p 355 p.

39. He X & Zhang J (2006) Toward a molecular understanding of pleiotropy. Genetics 173(4):1885–1891.

40. Flores CO, Poisot T, Valverde S, & Weitz JS (2016) BiMat: a MATLAB package to facilitate the analysis of bipartite networks. Methods Ecol Evol 7(1):127–132.

41. Wang Z & Zhang JZ (2007) In search of the biological significance of modular structures in protein networks. Plos Computational Biology 3(6):1011–1021.

42. Fritzemeier CJ, Hartleb D, Szappanos B, Papp B, & Lercher MJ (2017) Erroneous energy-generating cycles in published genome scale metabolic networks: Identification and removal. PLoS Comput Biol 13(4):e1005494.

43. Cameron FJ & Sinclair AH (1997) Mutations in SRY and SOX9: Testis-determining genes. Human Mutation 9(5):388–395.

44. McKusick-Nathans Institute of Genetic MedicineJHU Baltimore, MD (Online Mendelian Inheritance in Man (OMIM), https://omim.org.

45. Shlomi T, et al. (2007) Systematic condition-dependent annotation of metabolic genes. Genome Research 17(11):1626–1633.

46. Szappanos B, et al. (2016) Adaptive evolution of complex innovations through stepwise metabolic niche expansion. Nat Commun 7:11607.

47. The R Foundation (The R Project for Statistical Computing, https://www.r-project.org.

48. Gelius-Dietrich G, Desouki AA, Fritzemeier CJ, & Lercher MJ (2013) Sybil--efficient constraint-based modelling in R. BMC Syst Biol 7:125.

